# LUZP1 and the tumour suppressor EPLIN are negative regulators of primary cilia formation

**DOI:** 10.1101/736389

**Authors:** João Gonçalves, Étienne Coyaud, Estelle M.N. Laurent, Brian Raught, Laurence Pelletier

## Abstract

Cilia/flagella are microtubule-based cellular projections with important sensory and motility functions. Their absence or malfunction is associated with a growing number of human diseases collectively referred to as ciliopathies. However, the fundamental mechanisms underpinning cilia biogenesis and functions remain only partly understood. Here, we show that LUZP1, and its interacting protein EPLIN, are negative regulators of primary cilia formation. LUZP1 is an actin-associated protein that localizes to both actin filaments and the centrosome/basal body. Like EPLIN, LUZP1 is an actin-stabilizing protein and likely regulates actin dynamics at the centrosome. Both proteins interact with ciliogenesis and cilia length regulators, and are potential players in the actin-dependent processes involved in centrosome to basal body conversion. Ciliogenesis deregulation caused by LUZP1 or EPLIN loss may contribute to the pathology of their associated diseases.

**SUMMARY:** Gonçalves et al. show that LUZP1 and its interactor EPLIN are negative regulators of ciliogenesis. LUZP1 is a novel actin-stabilizing protein localizing at the centrosome and basal body where it may regulate actin-associated processes.

## INTRODUCTION

Eukaryotic cilia are microtubule (MT)-based organelles that protrude from the cell surface. In vertebrates, multiple immotile (i.e. primary cilia) and motile cilia fulfil critical sensory and motility functions required for embryonic development and in adult tissues homeostasis (Goetz and Anderson, 2010; Mirvis et al., 2018; Mitchell, 2007). Defects in cilia biogenesis and functions cause human diseases typified by symptoms like blindness, infertility, cognitive impairment and cystic kidneys (Mitchison and Valente, 2017). The ciliogenesis pathways are not fully known, but involve the cytoskeleton systems and multiple machineries including the ones participating in membrane trafficking, as well as centriolar satellites, which transport centrosomal/ciliary cargo to the centrosome and ciliary base (Hsiao et al., 2012; Mirvis et al., 2018; Odabasi et al., 2019). The MT and actin cytoskeletons act jointly in processes like cell adhesion, migration, centrosome positioning and mitotic spindle orientation (Dogterom and Koenderink, 2019), and also in ciliogenesis (Mirvis et al., 2018; Pitaval et al., 2017). Primary cilia biogenesis initiates at the centrosome, a MT and actin organizing center (Farina et al., 2016), and relies on its older (mother) centriole to become the basal body from which the ciliary axoneme is nucleated (Lu et al., 2015). During the formation of these cilia, a ciliary vesicle is formed at the distal end of the mother centriole/basal body, which then moves to the cell surface where it becomes attached to the cell membrane through structures called transition fibers. This migration process relies both on increased MT polymerization at the centrosome, as well as increased actin contractility (Pitaval et al., 2017). Interestingly, loss of function of actin regulators, as well as pharmacological disruption of the actin cytoskeleton (*e.g.* cytochalasin D and Jasplakinolide treatments) increases ciliation, and affects ciliary length and signaling in human cells (Kim et al., 2010; Nagai and Mizuno, 2017). Indeed, treating cells with low cytochalasin D doses leads to the accumulation of MyosinVa at the centrosome, which promotes the formation of the ciliary vesicle (Lu et al., 2015; Wu et al., 2018). Still, how global or centrosomal actin dynamics impact cilia biogenesis is not fully understood.

Here we show that LUZP1 is as an actin-associated protein which localizes to the centrosome, the basal body, actin filaments and the midbody. Moreover, it acts as a negative regulator of ciliogenesis, as its loss of function increases ciliation frequency in human RPE-1 (retinal pigmented epithelium) cells. Using proximity-dependent biotin identification (BioID) (Roux et al., 2012), co-immunoprecipitation (co-IP) and functional assays, we demonstrate that the actin stabilizing protein EPLIN interacts with LUZP1 and also negatively regulates ciliogenesis. Finally, we provide evidence supporting a role for LUZP1 in actin stabilization. These results are particularly relevant, as they may shed light on the molecular mechanisms of the disease LUZP1 and EPLIN are associated with, the 1p36 deletion syndrome (Jordan et al., 2015) and cancer (Collins et al., 2018; Jiang et al., 2008; Maul and Chang, 1999; Sanders et al., 2010; Zhang et al., 2011), respectively.

## RESULTS AND DISCUSSION

To address the fundamental question of primary cilia formation, we used BioID to characterize the interactome of key centrosomal and ciliary proteins in non-ciliated and ciliated human cells (Gupta et al., 2015). This allowed us detect alterations in the interactome upon ciliation that could illuminate potentially important events in the ciliogenesis programme. In this study, LUZP1 was identified as a prey for proteins that localize to centriolar satellites, the centrosome, and primary cilia, suggesting a centrosomal/ciliary localization and function for this protein (Gupta et al., 2015).

LUZP1 is a poorly characterized protein presenting an LCD1 (present in a family of proteins that regulate checkpoint kinases) (Rouse and Jackson, 2000) and three leucine zipper domains at its N-terminal region (Fig. 1 A) (Sun et al., 1996). LUZP1 is predominantly expressed in the mouse adult brain and neural lineages during development (Lee et al., 2001; Sun et al., 1996). To determine the subcellular localization of human LUZP1, we expressed it fused to GFP at the N- or C-terminus in RPE-1 cells, and performed immunofluorescence (IF) analyses with antibodies specific to GFP, and centrosomal and ciliary markers (Fig. 1 B, C; Fig. S1 A, B). Consistent with our proteomics data (Gupta et al., 2015), the fusion proteins accumulated at the centrosome and basal body of primary cilia. Also, they localized to filaments along the cytoplasm, which we confirmed to be actin fibers through fluorophore-labelled phalloidin co-staining (Figure 1C, D). Finally, LUZP1 was detected at the midbody during cytokinesis (Fig. 1 D). These results were supported by time-lapse imaging of RPE-1 cells stably expressing GFP-LUZP1, which showed the protein at the centrosome throughout the cell cycle, and strongly accumulating at the midbody during cytokinesis (Sup. Video 1). By treating RPE-1 cells expressing GFP-LUZP1 with nocodazole, we observed that the protein could still localize to the centrosome, suggesting that its localization there is MT-independent. (Fig. S1 C). To further refine the localization of LUZP1 at the ciliary base, we stained RPE-1 cells expressing GFP-LUZP1 with antibodies against proteins specific to different centriole and ciliary domains including centriolar sub-distal appendages (CEP170), distal appendages/transition fibers (CEP164) and the transition zone (CEP290) (Fig. S1 B). Our analysis revealed that LUZP1 does not localize to transition fibers or the transition zone but rather resides in the vicinity of sub-distal appendages, at the proximal end of basal bodies. To determine which protein domains within LUZP1 are important for its localization, we generated two truncation mutants (Fig. 1 A), one spanning only of the domains present at the N-terminus (aa 1-296), the other the remainder of the protein (aa 297-1076). The expression of these GFP-tagged mutants revealed that, like the full-length protein, the protein fragment lacking the leucine zipper and LCD1 domains localized to the centrosome, the basal body and actin filaments, showing they are not required for LUZP1 localization (Fig. 1 C).

**Figure 1.**
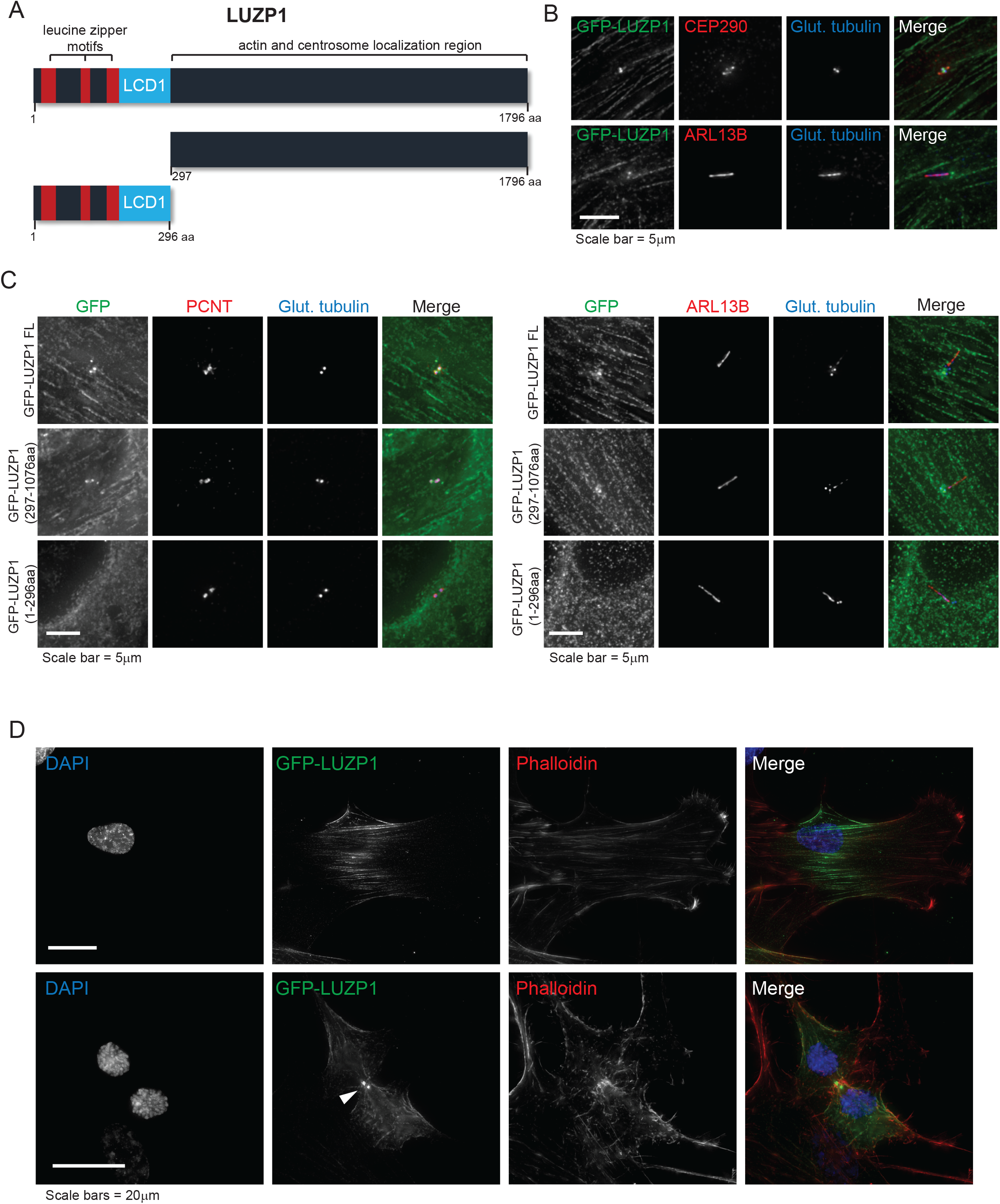
Sub-cellular localization of LUZP1, a novel centrosome/basal body and actin-associated protein. **A)** Schematic representation of human LUZP1, a 1796aa protein containing three leucine-zipper motifs and a LCD1 domain at its N-terminal region. Represented are also two truncation mutants. **B)** LUZP1 localizes to the centrosome and the basal body. RPE-1 cells stably expressing GFP-LUZP1 were stained for GFP, glutamylated tubulin (centrosome and ciliary marker) and CEP290 (centrosome and transition zone marker; top panel) or ARL13B (ciliary marker, bottom panel). **C**) The N-terminal region of LUZP1 is not required for the protein’s localization. RPE-1 stable cells stably expressing GFP-tagged LUZP1, full-length and the two truncation mutants represented in (A) were stained for GFP, glutamylated tubulin, and PCNT (centrosome marker; left panels) or ARL13B (right panels). **D)** LUZP1 localizes to actin filaments and the midbody (indicated by arrowhead). RPE-1 GFP-LUZP1 cells were stained using an antibody against GFP. The actin cytoskeleton was stained with fluorophore-conjugated phalloidin, and the DNA was stained with DAPI.

To assess the molecular pathways in which LUZP1 is involved, we defined its proximity interactome by performing BioID on a inducible Flp-In TREx 293 cell line to express FLAG-BirA*-tagged LUZP1 (Fig. S1 D). BirA* is an abortive mutant of an *E. coli* biotin ligase (R118G) which activates biotin and releases highly reactive biotinoyl-AMP in its vicinity (within a radius of ~10nm) (Roux et al., 2012). Epsilon amine groups of proximal proteins of FLAG-BirA*-LUZP1 get covalently conjugated with biotin. This biotin proximity labelled proteins are then captured by streptavidin-biotin affinity purification and identified by mass spectrometry (MS) (Roux et al., 2012). We showed that FLAG-BirA*-LUZP1 localizes to the centrosome and the actin cytoskeleton and we detected biotinylation signal in the same localizations using fluorophore-conjugated streptavidin (Fig. S1 D). BioID was performed both in cycling cells, and cells that were serum-starved to promote primary cilia formation. After background subtraction, we identified high confidence proximal interactors (Table 1; Sup. Table 1) that agree with our sub-cellular localization analysis and previous BioID studies. Indeed, except for EFHD1 and TP53BP2, the identified LUZP1 preys localize to centriolar satellites, the centrosome/cilium, and the MT and actin cytoskeletons (Table 1; Sup. Table 1), suggesting that LUZP1 has cytoskeleton-related roles. Previously, we identified LUZP1 as a proximity interactor of both PCM1 and CEP170 (Gupta et al., 2015). PCM1 is centriolar satellite component likely biotinylated by FLAG-BirA*-LUZP1 at the centrosome, since none of the LUZP1 fusions we tested localized to satellite granules (data not shown). Interestingly, among other proteins, PCM1 mediates the localization of actin regulators (*e.g.* Arp2/3 complex and WASH) to the centrosome, having an impact on this organelle’s role as an actin organizing center (Farina et al., 2016). CEP170 is a MT-binding protein present at centriolar sub-distal appendages/basal feet, that is required for the recruitment of MT and actin-associated ciliogenesis regulators to the mother centriole/basal body (Copeland et al., 2018). Another centrosome-related prey, identified only in cycling cells was MAP7D3, a MT-binding protein that regulates MT assembly and stability, being important for MT dynamics at spindle poles (Kwon et al., 2016; Sun et al., 2011; Yadav et al., 2014). Under serum starvation, we identified MRE11A and PPP1CA. MRE11A is a nuclease involved in DNA double–strand break repair that localizes to centrosomes (Zhao et al., 2008). PPP1CA is a phosphatase involved in MT and actin cytoskeleton processes including centrosome separation during cell division (Mi et al., 2007). Two molecular chaperones were identified as LUZP1 interactors. HSPA1B, a member of the heat shock protein 70 family, localizes to mitotic centrosomes being important for their maturation by facilitating the recruitment of pericentriolar material components (Fang et al., 2019). Interestingly, this protein was only detected in cycling cells possibly due to the reduced number of mitotic cells upon serum-starvation. On the other hand, CCT3, a subunit of the chaperonin containing TCP-1 (CCT), was detected under serum-starvation conditions. CCT is a molecular chaperone involved in the folding of tubulin, actin and hundreds of other proteins. CCT subunits were also proposed to have roles outside the chaperonin complex, and some localize at centrosomes and cilia (Goncalves et al., 2010; Seixas et al., 2003; Seixas et al., 2010; Sinha et al., 2014) where they may play roles related to the MT and actin-related processes of ciliogenesis. Myosin subunits were also identified, some of which (MYL12A/B, MYL3 or MYL6/B) only in serum-starved cells. Moreover, DAPK3, which regulates myosin by phosphorylating MYL9 and MYL12B (Komatsu and Ikebe, 2004; Murata-Hori et al., 1999), was also detected. Myosins are actin-binding molecular motors that play several roles in processes such as cell migration and division (Hartman and Spudich, 2012). Of note, myosins can accumulate at the centrosome and primary cilium upon actin destabilization (Wu et al., 2018). Therefore, our results may reflect actin cytoskeleton changes happening during ciliogenesis. Another protein identified in the LUZP1 BioID interactome only in serum-starved cells was SEPT2. Septins interact with the MT and actin cytoskeletons and exert scaffolding roles. SEPT2 forms a complex with SEPT7 and SEPT9 which localizes along the axoneme of primary cilia and is required for ciliogenesis (Ghossoub et al., 2013; Hu et al., 2010). Four other proteins with actin cytoskeleton-associated roles were also found in our study. TJP1 is part of the tight junction structure, which interacts with actin to establish cellular connections (Fanning et al., 1998). SORBS1/CAP, is a multidomain protein that binds to several cytoskeletal proteins, and plays a role in the formation of actin stress fibers and focal adhesions (Ribon et al., 1998). In agreement, it regulates processes like cell adhesion, spreading and motility (Zhang et al., 2006). Filamins (FLNA/B) have important actin-related functions by binding the actin cytoskeleton network to membrane components (Nakamura et al., 2011). Of note, FLNA interacts with the ciliary protein MKS3, and this interaction is important during ciliogenesis for the correct positioning of the basal body (Adams et al., 2012). LUZP1 was recently identified as a FLNA binding partner, further supporting our BioID results (Wang and Nakamura, 2019). Finally, EPLIN/LIMA1 binds to, cross-links and stabilizes actin filaments (Maul et al., 2003). In addition, it plays important roles at cellular junctions and focal adhesions (Abe and Takeichi, 2008; Karakose et al., 2015). LUZP1 was identified as an EPLIN interactor in a previous proteomics study that used affinity purification followed by MS to identify the interactome of over 1000 human proteins in HeLa cells (Hein et al., 2015). Having two distinct MS approaches identifying the same interaction in two cell lines strongly supports EPLIN and LUZP1 associate and are functionally related. Also, both LUZP1 and EPLIN are part of the BioID interactome of the human phosphatase CDC14A, which modulates actin through dephosphorylation of EPLIN (Chen et al., 2017), and regulates actin nucleation at the centrosome impacting primary cilia length (Uddin et al., 2019). Taking this into account, we decided to prioritize the validation of the LUZP1-EPLIN interaction and study the role of both proteins in parallel.

**Table 1.**
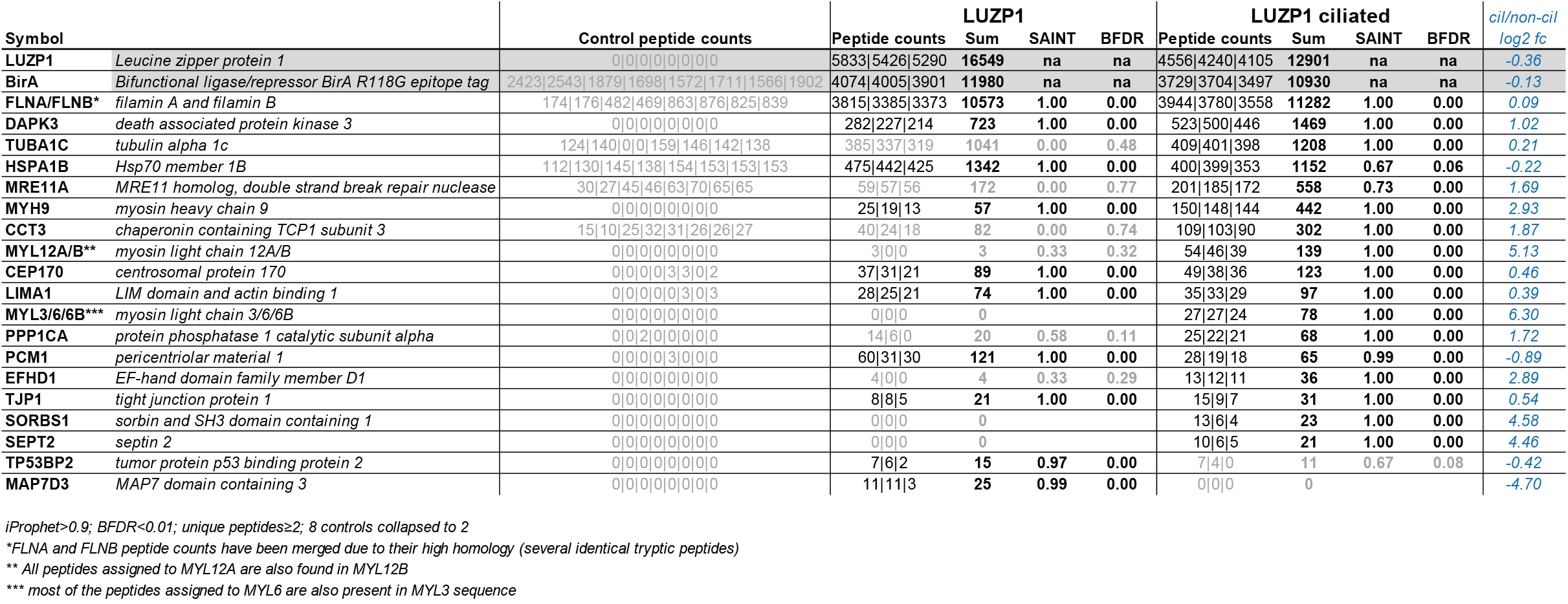

Two alternative promoters drive the expression of the *LIMA1* gene resulting in the expression of EPLIN isoforms α and β (Fig. 2 A) (Chen et al., 2000). Both isoforms contain a central LIM domain possibly involved in protein-protein interactions (Maul and Chang, 1999). Also, EPLIN proteins have at least two actin binding domains (Maul et al., 2003). The two isoforms are differently expressed in a number of cell lines and tissues (Maul and Chang, 1999; Maul et al., 2001; Wang et al., 2007), with RPE-1 cells expressing mostly EPLINα (Fig. 2 B). To study the localization of EPLIN, and compare it to that of LUZP1, we generated RPE-1 cells expressing GFP-tagged EPLIN isoforms alone, or in combination with mCherry-LUZP1 (Fig. 2 C; Fig. S2 A). Both EPLIN isoforms localized to actin filaments, however, isoform α tended to accumulate on filopodia and membrane ruffles at the cell cortex, often at what was clearly the leading edge of a moving cell. In contrast to LUZP1, none of the EPLIN isoforms was observed at the centrosome, basal body, or midbody (Fig. 2 C; Fig. S2 A, B).

**Figure 2.**
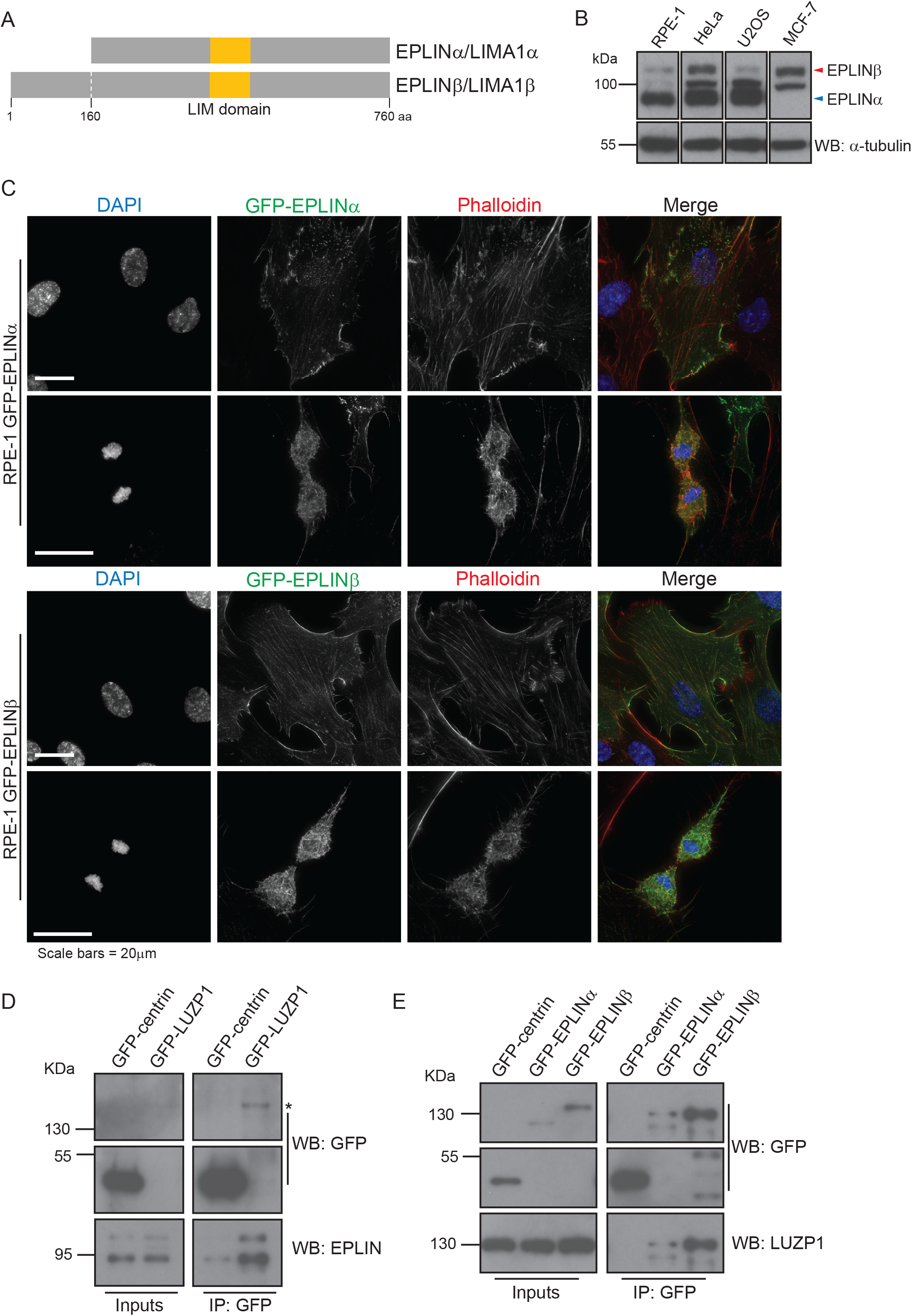
EPLINα/β interacts with LUZP1. **A)** Schematic representation of human EPLIN isoforms α (600 aa) and β (760 aa) both of which contain a LIM domain. **B)** EPLIN isoforms are expressed differently in distinct cell lines. Western blot analysis of EPLIN expression in human RPE-1, HeLa, U2OS and MCF-7 cells. **C)** EPLIN localizes to actin structures in RPE-1 cells. RPE-1 cells stably expressing GFP-EPLINα or β cells were stained with an antibody against GFP. The actin cytoskeleton was stained with fluorophore-conjugated phalloidin, and the DNA was stained with DAPI. **D**) GFP-LUZP1 pulls-down endogenous EPLIN. Co-immunoprecipitation experiments using protein extracts prepared from RPE-1 cells stably expressing GFP-centrin (control) or GFP-LUZP1. The fusion proteins were immunoprecipated using GFP antibody-conjugated beads. GFP and EPLIN antibodies were used to detect the GFP fusions and endogenous EPLIN isoforms, respectively. * indicates the GFP-LUZP1 band. **E)** Both EPLIN isoforms pull-down endogenous LUZP1. Co-immunoprecipitation experiments using protein extracts prepared from RPE-1 cells stably expressing GFP-centrin (control) or GFP-EPLINα/β. The fusion proteins were immunoprecipated using GFP antibody-conjugated beads. GFP and LUZP1 antibodies were used to detect the GFP fusions and endogenous LUZP1, respectively.

To validate the interaction between LUZP1 and EPLIN, we performed co-IP experiments using the RPE-1 stable lines, and observed that GFP-tagged LUZP1 pulled-down both endogenous EPLIN isoforms (Fig. 2 D). Similarly, GFP-tagged EPLINα and β pulled-down endogenous LUZP1 (Fig. 2 E). We further showed that FLAG-tagged EPLINβ pulled-down endogenous LUZP1 in HEK293 cells, indicating the observed interactions were not dependent on the tag used for the co-IPs (Fig. S2 C). To determine if LUZP1 leucine zipper and LCD1 domains are important for the interaction with EPLIN, we expressed GFP-EPLINβ in HEK293 stable/inducible lines expressing different constructs: FLAG-LUZP1 (full length), FLAG-LUZP1 (aa 1-296), or FLAG-LUZP1 (aa 297-1076). By co-IP, we observed that GFP-EPLINβ pulled-down only full-length FLAG-LUZP1 and FLAG-LUZP1 (aa 297-1076) showing that the C-terminal region of LUZP1 mediates the interaction with EPLIN (Fig. S2 D). Finally, we also showed that FLAG-LUZP1, and FLAG-EPLINβ, pull-down actin (Fig. S2 C, E).

Next, we investigated the role of LUZP1 and EPLIN by determining their loss of function phenotypes in RPE-1 cells. Previously, LUZP1 scored as a negative regulator of ciliogenesis in an RNAi screen of the centrosome-cilium interface proximity interactome (Gupta et al., 2015). Taking this into account, as well as LUZP1 localization to the basal body, we decided to investigate a potential role for this protein in ciliogenesis by conducting RNAi experiments. After transfection with a negative control siRNA or siRNAs targeting LUZP1, the cells were serum-starved for 72h to induce primary cilia formation. In agreement with the RNAi screen, we observed a higher number of ciliated cells in the LUZP1 RNAi condition (Fig. S3 A). To explore these results further, and confirm the specificity of the phenotype, we tested if LUZP1 depletion could induce ciliogenesis in conditions that do not favor it, *i.e*. in the presence of serum. We also tested if the phenotype is specific by using RPE-1 cells expressing GFP-LUZP1 constructs, one RNAi-sensitive, the other resistant. As a control, we used cells expressing GFP only. The different cell lines were transfected with control or LUZP1 siRNAs for 72h, but were not serum-starved. In these conditions, we observed a significant increase in the number of ciliated cells in the LUZP1-depleted populations, except in the cell line expressing the RNAi-resistant construct (Fig. 3 A, B). Western blot analysis confirmed the depletion of endogenous and GFP-tagged LUZP1 in the sensitive cell line, but only of the endogenous protein in the resistant one (Fig. 3 C). These results show that the phenotype is specific, and confirm LUZP1 as a negative regulator of primary cilia formation.

**Figure 3.**
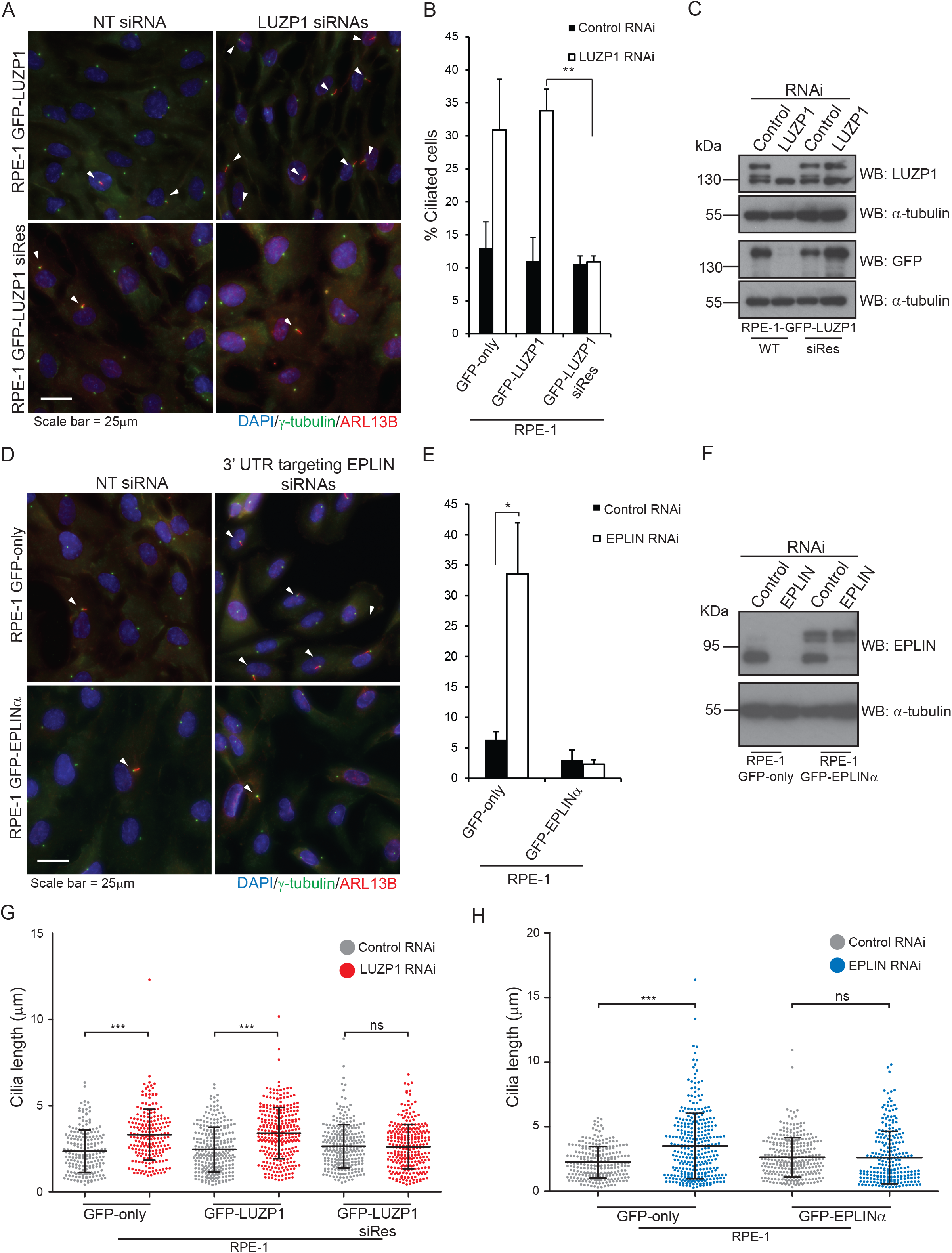
LUZP1 and EPLIN are novel negative regulators of ciliogenesis. **A)** LUZP1 depletion increases ciliation in RPE-1 cells. Immunofluorescence analysis of RPE-1 cells stably expressing siRNA sensitive or resistant GFP-LUZP1 and transfected with control (NTsiRNA) or siRNAs targeting LUZP1 for 72h. The cells were stained for γ-tubulin (centrosome and basal body marker) and ARL13B (ciliary marker). DNA was stained with DAPI. The arrowheads indicate primary cilia. **B)** Bar graph shows the mean percentage of ciliated cells (n > 200 cells per sample, 3 independent experiments) in both RPE-1 stable lines transfected with the indicated siRNAs for 72h. Error bars indicate SD. **p < 0.01 (Student’s two-tailed t test). **C)** Western blot analysis of endogenous and GFP-tagged LUZP1 in both stable lines transfected with control non-targeting (NT), or LUZP1-directed siRNAs for 72h. **D)** EPLIN silencing increases ciliation in RPE-1 cells. Immunofluorescence analysis of RPE-1 cells stably expressing GFP-only or GFP-EPLINα and transfected with control (NT siRNA) or siRNAs targeting the 3’ UTR of EPLIN for 72h. The cells were stained for γ-tubulin (centrosome and basal body marker) and ARL13B (ciliary marker). DNA was stained with DAPI. The arrowheads indicate primary cilia. **E)** Bar graph shows the mean percentage of ciliated cells (n > 200 cells per sample, 3 independent experiments) in both stable lines transfected with the indicated siRNAs for 72h. Error bars indicate SD. *p < 0.05 (Student’s two-tailed t test). **F)** Western blot analysis of endogenous and GFP-tagged EPLINα in the cells transfected with control non-targeting (NT), or LUZP1-directed siRNAs for 72h. **G, H)** LUZP1 and EPLIN depletion leads to increased ciliary length in RPE-1 cells. The graphs show the length of cilia measured in the indicated populations treated wither with control siRNA or siRNAs directed against LUZP1 or EPLIN. n> 200 cilia per condition (data pooled from three independent experiments). The bars correspond to the average and SD. ***p < 0.001 (Student’s two-tailed t test); ns – non significant.

Next, we tested if a similar phenotype would be observed for EPLIN. For this assay, RPE-1 cells expressing GFP only or GFP-EPLINα were transfected with control siRNAs or siRNAs targeting the 3’-UTR of *EPLIN* transcripts. The siRNAs efficiently depleted endogenous EPLIN but not the GFP fusion (Fig. 3 F). 72h post-transfection in the presence of serum, we observed a significant increase in the number of ciliated cells in the control cell line, but not in the one expressing GFP-EPLINα (Fig. 3 D-F). Thus, like LUZP1, EPLIN is a negative regulator of ciliogenesis. To rule out cell cycle arrest as a cause for the increased ciliation in cells depleted of LUZP1 and EPLIN, we followed their growth throughout the experiment by time-lapse imaging, and showed that cell proliferation was not perturbed by the RNAi of any of the genes (Sup. Fig. 3B). Given the known function of EPLIN as an actin stabilizer (Maul et al., 2003), LUZP1 localization to the centrosome/basal body and actin filaments, and actin-associated BioID preys, we hypothesized that affected actin-related processes might be causing the increased ciliation. Indeed, disrupting the actin cytoskeleton pharmacologically or through the depletion of actin regulators causes increased ciliation and primary cilia lengthening (Kim et al., 2010). Hence, we tested if ciliary length was affected in cells depleted of LUZP1 and EPLIN. By measuring cilia length in the cell populations described above, we showed that LUZP1 or EPLIN-silenced cells had significantly longer primary cilia than control cells. These phenotypes were rescued by the expression of the respective protein as a GFP-fusion, confirming their specificity (Fig. 3 G, H). Given the similarity in loss of function phenotypes between LUZP1 and EPLIN we posited that LUZP1 could also stabilize the actin cytoskeleton. To test this, we used an approach that supported EPLIN’s actin stabilization properties (Maul et al., 2003). Specifically, we transiently expressed FLAG-tagged LUZP1 or EPLIN in RPE-1 and MCF-7 cells, and stained them for actin. Compared to control cells transfected with the empty plasmid, and non-expressing neighboring cells, cells over-expressing FLAG-LUZP1 or FLAG-EPLIN showed brighter actin filaments, suggesting their stabilization (Fig. 4 A; Fig. S3C). Also, in agreement with a recent report showing that LUZP1 fragments (aa 1-500; aa 400-500) cross-link actin-filaments *in vitro* (Wang and Nakamura, 2019), the over-expression of FLAG-tagged LUZP1 aa 1-496 in RPE-1 cells dramatically affected actin organization, with the filaments appearing curved likely due to bundling (Fig. 4 B). As previously mentioned, low doses of cytochalasin D, an inhibitor of actin polymerization, induce ciliation. To further investigate LUZP1 potential actin stabilization role, we posited that over-expressing it could counteract the effect of cytochalasin D on ciliation. To test this hypothesis, we used two control cell lines (wild-type and GFP-only RPE-1 cells), and cell lines expressing GFP-LUZP1, GFP-EPLINα, GFP-EPLINβ, or the EPLIN fusions in combination with mCherry-LUZP1. The cells were treated with a low cytochalasin D dose (50nM) for 16h in the presence of serum, after which ciliation was assessed. As expected, this caused a significant increase in the number of ciliated cells in control cells. However, in the over-expressing lines the increase in ciliation was significantly less (~ half) than in controls. Interestingly, over-expressing both LUZP1 and EPLIN did not have an additive effect on ciliation prevention (Fig. S3 D).

**Figure 4.**
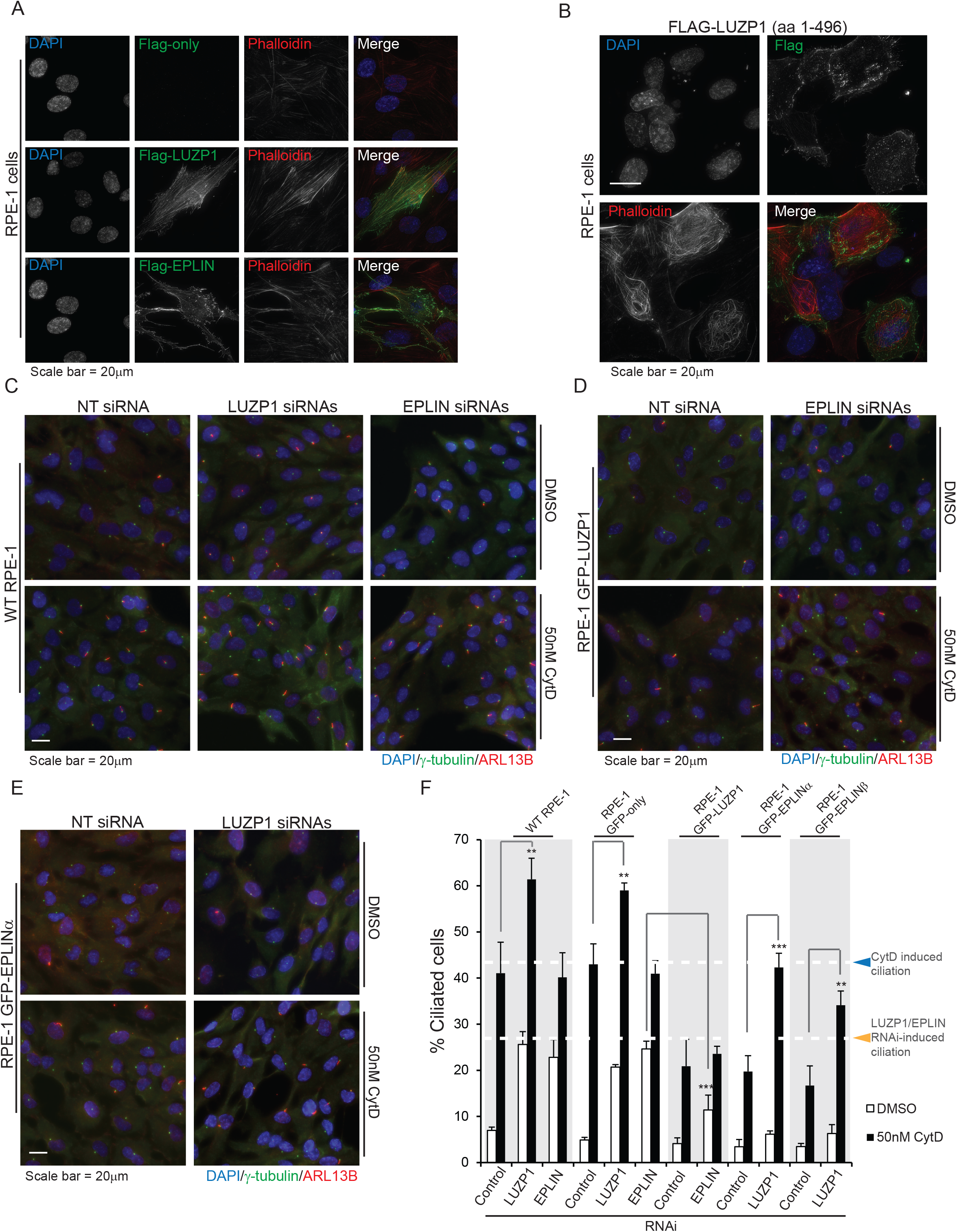
LUZP1 and EPLIN have actin stabilization roles. **A)** Immunofluorescence analysis of RPE-1 cells transiently transfected with a control empty Flag vector or plasmids to express FLAG-LUZP1 or FLAG-EPLINβ. The fusion proteins were detected with a FLAG antibody. The actin cytoskeleton was stained with fluorophore-conjugated phalloidin, and the DNA was stained with DAPI. **B)** Immunofluorescence analysis of RPE-1 cells transiently transfected with a FLAG-LUZP1 (aa 1-496) construct. The actin cytoskeleton was stained with fluorophore-conjugated phalloidin, and the DNA was stained with DAPI. **C-E)** Wild-type and transgenic RPE-1 cells were transfected for 72h with control siRNA or siRNAs targeting LUZP or EPLIN, as indicated. During the last 16h of transfection, the cells were treated either with DMSO (control) or 50nM cytochalasin D. The cells were stained for γ-tubulin (centrosome and basal body marker) and ARL13B (ciliary marker). DNA was stained with DAPI. **F)** Bar graph shows the mean percentage of ciliated cells (n > 200 cells per sample, 3 independent experiments) in the indicated cell lines, transfected with control, LUZPI or EPLIN siRNAs and treated with DMSO or 50nM cytochalasin D. Error bars indicate SD. **p < 0.01; ***p < 0.001 (Student’s two-tailed t test).

Collectively, our results so far suggested that LUZP1 and EPLIN have similar functions regarding ciliogenesis regulation, possibly through their actin-associated roles. Yet, it remained unclear if they act in the same pathways. To test this, we used RNAi to simultaneously deplete LUZP1 and EPLIN in wild-type RPE-1 cells, and assessed the effect of their depletion on cilia formation in the presence of serum. Our results showed that co-depleting these proteins caused a significant increase in ciliation compared to the individual RNAi conditions (Fig. S3 E, F), suggesting LUZP1 and EPLIN may not fully overlap in the molecular mechanisms they are involved in. To further explore the role of both proteins, we tested the effect of cytochalasin D treatment combined with the RNAi of LUZP1 or EPLIN on ciliogenesis. For this, we used two control cell lines (wild-type and GFP-only RPE-1 cells), and the lines expressing GFP-tagged LUZP1, EPLINα or EPLINβ (Fig. 4 C-F; Fig. S3 G, H). Cells were transfected for 72h in the presence of serum, and were treated with cytochalasin D or DMSO (vehicle) for the last 16h. The control cell lines treated with DMSO, showed the expected increase in ciliation caused by LUZP1 or EPLIN depletion (Fig. 4 C, F). Also as expected, the treatment with cytochalasin D increased ciliation in the control cell lines transfected with non-targeting siRNAs. Strikingly, the same cell lines depleted of LUZP1, but not EPLIN, showed an additional significant increase in the percentage of ciliated cells upon treatment with the drug (Fig. 4 C, F). One possible explanation for these results is that LUZP1 depletion destabilizes the actin cytoskeleton, sensitizing the cells to this actin-disrupting drug. This may be particularly relevant at the centrosome where LUZP1 could have an important role in modulating actin dynamics during the centrosome to basal body transition. Instead, LUZP1 may have other functions that account for its role in ciliogenesis, and what we observed is the additive effect of its depletion and the drug effect. Interestingly, silencing EPLIN did not produce the same effect in combination with cytochalasin D (Fig. 4 C, F), which may suggest it acts in the same pathway affected by the drug. In RPE-1 GFP-LUZP1 cells treated with DMSO and depleted of EPLIN, ciliation increased to a lesser extent than in the control cell lines in the same condition (Fig. 4 D, F). This suggests that over-expressing LUZP1 can rescue the effect of EPLIN depletion in cilia formation at least partially. In agreement with the aforementioned results (Fig. S3 D), treatment with cytochalasin D lead to a significantly lower level of induced ciliation in these cells. Also, EPLIN depletion and cytochalasin D treatment didn’t have a synergistic effect in increasing cilia formation (Fig. 4 D, F). Finally, in GFP-EPLINα or β expressing cells treated with DMSO, LUZP1 depletion did not lead to a significant increase in ciliation suggesting that the actin-stabilizing effect of over-expressed EPLIN rescues the LUZP1 RNAi phenotype (Fig. 4 E, F; Fig. S3 G). Upon cytochalasin D treatment, EPLIN over-expressing cells transfected with control siRNAs showed the expected modest increase in ciliogenesis. However, the same cells depleted of LUZP1 and treated with cytochalasin D showed ciliation levels similar to those of the control cell lines treated with the drug (Fig. 4 E, F; Fig. S3 G). This suggests that EPLIN requires LUZP1 to counteract the actin disruption and ciliation increase caused by cytochalasin D. These results may be related to the centrosome localization of LUZP1 and actin dynamics at this organelle. Indeed, centrosomal LUZP1 might be necessary to translate the actin stabilization effect of over-expressed EPLIN to the centrosome in extreme conditions of compromised actin functions such as the treatment with cytochalasin D. Further work will be necessary to determine the role(s) of LUZP1, EPLIN, and their interactors, on the local actin-related processes involved in centrosome to basal body transition and primary cilia formation.

Here we describe two new players in the ciliogenesis pathways. We showed that human LUZP1 localizes to the centrosome, the basal body, actin filaments and the midbody, which agrees with its interactome, composed mainly of centrosome/basal body components, and actin-associated proteins like EPLIN. LUZP1 depletion in RPE-1 cells did not affect cell cycle progression. In fact, despite the midbody localization, no signs of cytokinesis failure (e.g. unresolved cellular bridges; multinucleated cells; abnormal centrosome numbers) in LUZP1-depleted cells were observed. Still, we cannot rule out a function for LUZP1 in cell division, as remaining levels of the protein after RNAi may be enough to fulfil it. Silencing LUZP1 and EPLIN caused increased ciliation and ciliary lengthening. These results and the similarity to EPLIN in terms of loss of function phenotypes, lead us to hypothesise an actin-related role for LUZP1, which was supported by several lines of evidence. First, similar to EPLIN, LUZP1 over-expression stabilizes actin as evidenced by brighter phalloidin staining of actin filaments. Also, over-expressing aa 1-496 caused a striking effect on actin organization consistent with a recent report showing that LUZP1 fragments (aa 1-500, and aa 400-500) cross-link actin *in vitro* (Wang and Nakamura, 2019). These results suggest that the C-terminal region of LUZP1 (aa 500-1796) somehow masks/inhibits this actin-bundling domain. Second, the over-expression of LUZP1, as well as that of EPLIN, significantly rescues the increased ciliation phenotype caused by cytochalasin D. Also, depleting LUZP1, but not EPLIN, sensitizes cells to cytochalasin D treatment, reflected by an additional increase in ciliation levels when the two treatments are combined. Of note, our findings that LUZP1 is a regulator of ciliogenesis and an actin-stabilizing protein have been validated in a recent report (Bozal-Basterra et al., 2019).

The question still remains regarding the relevance of the interaction between LUZP1 and EPLIN. There are similarities in loss of function and over-expression phenotypes for both proteins. Also, the over-expression of one can rescue, at least partially, the loss of function phenotype of the other. However, the co-depletion of both genes leads to an additional increase in ciliation compared to individual knockdowns. Importantly, LUZP1 is required for EPLIN’s role in counteracting the effect of cytochalasin D on ciliation. One key difference between the two proteins relates to their localization. LUZP1, but not EPLIN, localizes to the centrosome/basal body where it may regulate actin dynamics. This could be critical to translate EPLIN’s effect on actin to the site of cilia formation in situations of a compromised cytoskeleton, as under cytochalasin D treatment. Alternatively, LUZP1 has other roles not necessarily actin-related. Understanding LUZP1 function is important, since its loss may contribute to the 1p36 deletion syndrome. Terminal deletions of chromosome 1p36 cause this developmental disease with potential clinical manifestations being short stature, affected brain development (*e.g.* microcephaly), hearing loss, congenital heart disease and renal disease (Zaveri et al., 2014). The phenotypes of a LUZP1 knockout mouse model (cardiovascular defects; cranial neural tube closure failure) agree with the hypothesis that the loss of the human gene contributes to this disorder. Importantly, the mice show ectopic sonic-hedgehog signaling, which may be due to perturbed ciliation (Hsu et al., 2008). On the other hand, EPLIN loss of expression has been associated with cancer. This has been shown to affect cancer cell adhesion and migration, and increase metastatic potential (Collins et al., 2018; Jiang et al., 2008; Liu et al., 2012; Sanders et al., 2010; Zhang et al., 2011). Like LUZP1, EPLIN had never been implicated in cilia biology. However, these proteins’ roles as negative regulators of ciliogenesis likely impact the multiple cilia-dependent signaling pathways contributing to the etiology of their associated diseases. Indeed, several of 1p36 deletion syndrome phenotypes are similar to those of well-established ciliopathies, and perturbed ciliary signaling contributes to the development of certain cancers.

## ABBREVIATIONS

BioID: Biotin identification
Co-IP: Co-immunoprecipitation
IF: Immunofluorescence
MT: Microtubule
MS: Mass spectrometry

---

**Supplementary Figure 1.**
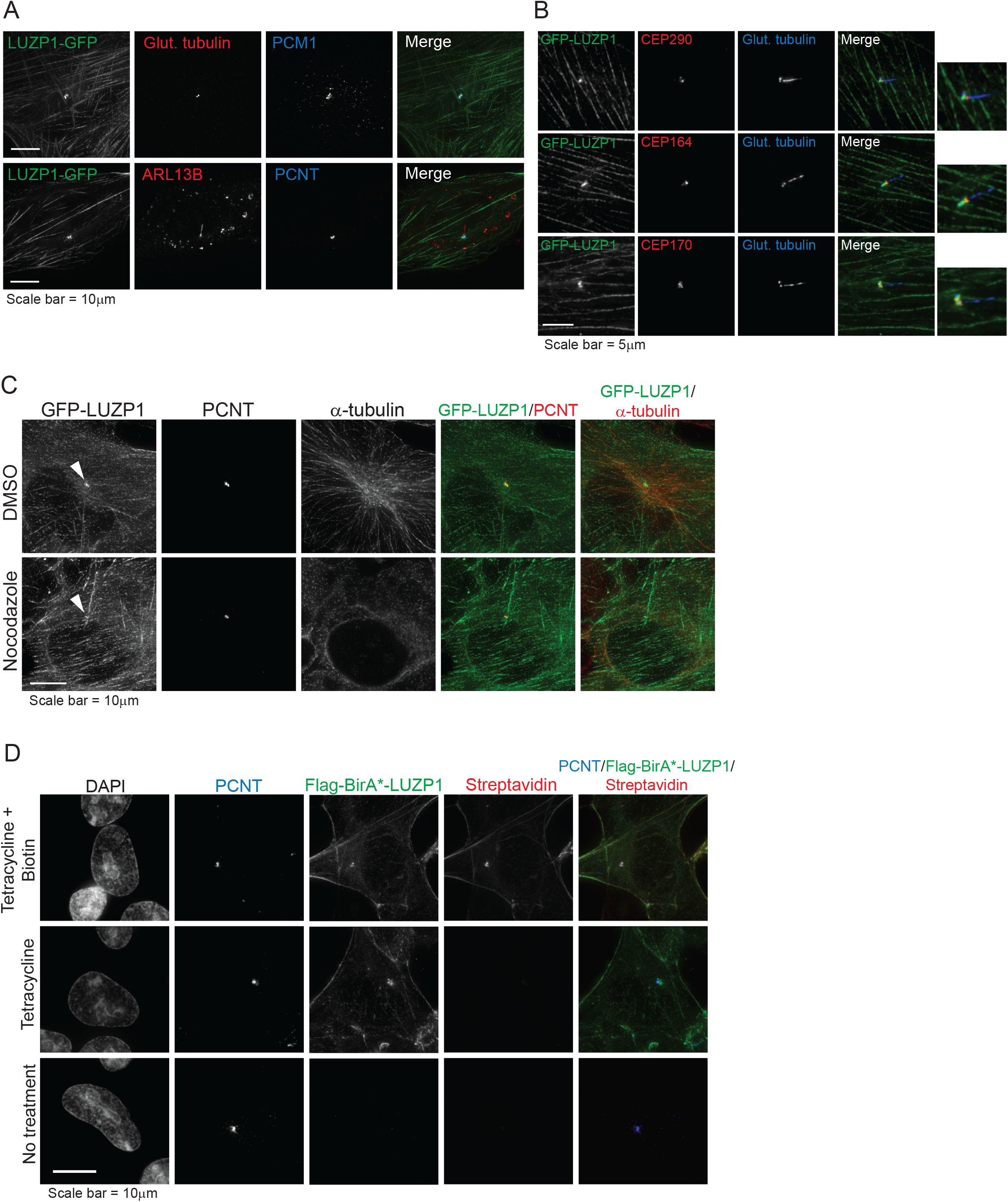
LUZP1 localizes to the proximal end of centrioles and its centrosomal localization is independent of microtubules. **A)** RPE-1 cells transiently expressing LUZP1-GFP cells were stained for GFP, glutamylated tubulin (centrosome and ciliary marker) and PCM1 (centriolar satellite marker; top panel), or ARL13B (ciliary marker) or PCNT (centrosome marker; bottom panel). **B)** RPE-1 cells stably expressing GFP-LUZP1 were stained for GFP, glutamylated tubulin (centrosome and ciliary marker) and CEP290 (centrosome and transition zone marker; top panel), CEP164 (distal appendage/transition fiber marker) or CEP170 (sub-distal appendage marker). **C)** Immunofluorescence analysis of RPE-1 GFP-LUZP1 stable cells treated with DMSO or Nocodazole. The cells were stained for GFP, PCNT and α-tubulin. The arrow head indicates the centrosome. **D)** Immunofluorescence analysis of HEK293 FLAG-BirA*-LUZP1 stable/inducible cells without treatment, treated with tetracycline only, or with tetracycline and biotin. The cells were stained with antibodies against FLAG and PCNT. Biotinylated proteins were detected with fluorophore-conjugated streptavidin and DNA with DAPI.

**Supplementary Figure 2.**
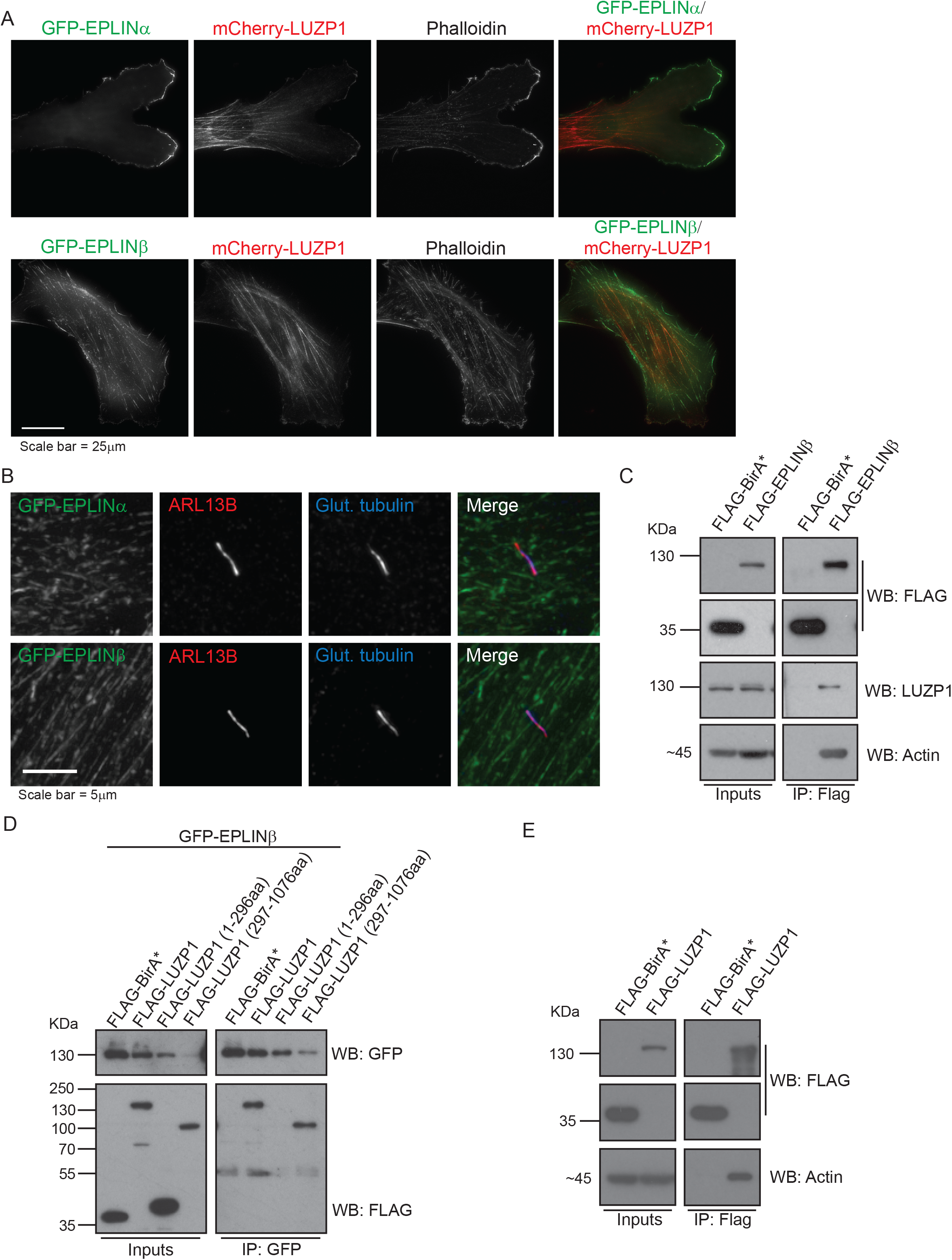
EPLIN sub-cellular localization and interaction with LUZP1. **A)** RPE-1 stable cells expressing GFP-EPLINα or β, and mCherry-LUZP1 were fixed and stained with phalloidin (actin market). EPLINα/β localize along actin filaments, and EPLINα accumulates at the cortex of the leading edge. **B)** Immunofluorescence analysis of RPE-1 stable cells expressing GFP-EPLINα or β were stained for ARL13B (ciliary marker) and glutamylated tubulin (centriole and ciliary marker). **C)** FLAG-EPLINβ pulls-down endogenous LUZP1. Co-immunoprecipitation experiments using protein extracts prepared from HEK293 cells stably expressing FLAG-BirA* (control) or FLAG-EPLINβ. The fusion proteins were immunoprecipated using FLAG antibody-conjugated beads. FLAG, LUZP1 and actin antibodies were used to detect the FLAG fusions, endogenous LUZP1 and actin, respectively. **D)** GFP-EPLINβ pulls-down the LUZP1 truncation mutant lacking the N-terminal domains. Co-immunoprecipitation experiments using protein extracts prepared from HEK293T stable cells expressing FLAG-BirA* (control), FLAG-LUZP1, FLAG-LUZP1 (1-296aa) or FLAG-LUZP1 (297-1076aa), and GFP-EPLINβ. The GFP-fusion was immunoprecipated using GFP antibody-conjugated beads. GFP and FLAG antibodies were used to detect the GFP and FLAG fusions, respectively. **E)** FLAG-LUZP1 pulls-down actin. Co-immunoprecipitation experiments using protein extracts prepared from HEK293 cells stably expressing FLAG-BirA* (control) or FLAG-LUZP1. The fusion proteins were immunoprecipated using FLAG antibody-conjugated beads. FLAG and actin antibodies were used to detect the FLAG fusions and endogenous actin, respectively.

**Supplementary Figure 3.**
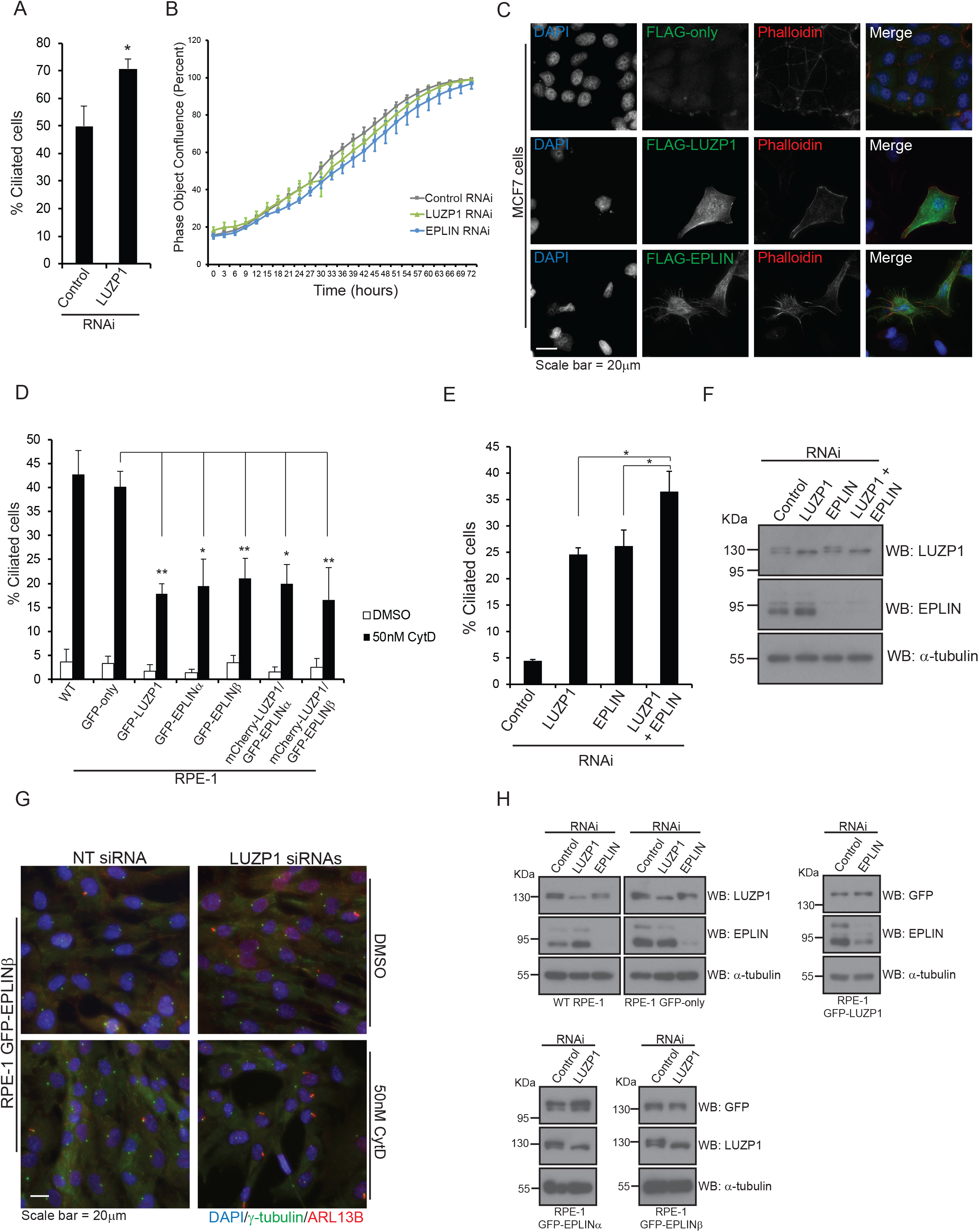
LUZP1 is an actin-stabilizing protein. **A)** LUZP1 depletion increases ciliation in RPE-1 cells. Wild-type RPE-1 cells were transfected with control (NTsiRNA) or siRNAs targeting LUZP1and serum-starved for 72h. Bar graph shows the mean percentage of ciliated cells (n > 200 cells per sample, 3 independent experiments) in RPE-1 cells transfected with the indicated siRNAs for 72h. Error bars indicate SD. *p < 0.05 (Student’s two-tailed t test). **B)** LUZP1 and EPLIN silencing does not affect cell proliferation. RPE-1 cells were transfected with a control siRNA or siRNAs targeting LUZP1 or EPLIN. Cell confluence was assessed by time-lapse imaging (determined by phase-contrast images). **C)** Immunofluorescence analysis of MCF7 cells transiently transfected with a control empty FLAG vector or plasmids to express FLAG-LUZP1 or FLAG-EPLINβ. The fusion proteins were detected with a Flag antibody. The actin cytoskeleton was stained with fluorophore-conjugated phalloidin, and the DNA was stained with DAPI. **D)** Bar graph shows the mean percentage of ciliated cells (n > 200 cells per sample, 3 independent experiments) in the indicated cell lines treated with DMSO or 50nM cytochalasin D. Error bars indicate SD. *p < 0.05; **p < 0.01 (Student’s two-tailed t test). **E)** Wild-type RPE-1 cells were transfected for 72h with a control siRNA, or siRNAs targeting LUZP1 or EPLIN, individually or in combination. Bar graph shows the mean percentage of ciliated cells (n > 200 cells per sample, 3 independent experiments) in each population. Error bars indicate SD. *p < 0.05 (Student’s two-tailed t test). **F)** Western blot analysis of endogenous and LUZP1 and EPLIN in RPE-1 cells transfected with the indicated siRNAs for 72h. **G)** Immunofluorescence analysis of RPE-1 GFP-EPLINβ cells transfected for 72h with control siRNA or siRNAs targeting LUZP. During the last 16h of transfection, the cells were treated either with DMSO (control) or 50nM cytochalasin D. The cells were stained for γ-tubulin (centrosome and basal body marker) and ARL13B (ciliary marker). DNA was stained with DAPI. **H)** Western blot analysis using protein extracts prepared from RPE-1 control cell lines and lines expressing GFP-LUZP1 or GFP-EPLINα or β. The cells were transfected with a control siRNA or siRNAs targeting LUZP1 or EPLIN, as indicated. A GFP antibody was used to detect the GFP-fusions and antibodies against LUZP1 and EPLIN were used to detect the respective endogenous proteins.

## Supplementary Video 1

**LUZP1 localizes to actin filaments, centrosomes and the midbody**

RPE-1 cells stably expressing GFP-LUZP were imaged every 5 min under controlled temperature and CO_2_ growth conditions. Arrow heads point at centrosomes in interphase and mitotic cells and the arrow points at the midbody. The video is played at 7 fps.

## MATERIAL AND METHODS

### Cell culture

Flp-In™ T-REx™ HEK293 cells were grown in Dulbecco’s Modified Eagle’s Medium (DMEM) supplemented with 10% fetal bovine serum (FBS), GlutaMAX™, zeocin (100μg/ml) and blasticidin (3μg/ml). Flp-In™ T-REx™ 293 stable lines expressing Flag-BirA* or Flag-BirA*-LUZP1 were maintained as above, with the addition of hygromycin (200μg/ml) or puromycin (1μg/ml). HeLa and MCF-7 cells were grown in DMEM supplemented with 10% FBS and GlutaMAX™. hTERT□RPE□1 cells (WT and stable lines) were grown in DMEM/F12 supplemented with 10% FBS, GlutaMAX™ and sodium bicarbonate (1.2 g/l). U-2 OS cells were grown in McCoy 5A medium supplemented with 10% FBS and GlutaMAX™. All cells were cultured in a 5% CO_2_ humidified atmosphere at 37 °C. hTERT□RPE□1 cells stably expressing GFP-centrin were a generous gift from Dr. A. Khodjakov.

### Molecular cloning

The human LUZP1 (NM_001142546.1) and EPLIN (NP_001107018.1; NM_001113547.2) coding sequences were amplified from a testes cDNA sample using primers (Supplementary Table 2) containing appropriate restriction sites. The PCR products were digested and ligated into the following vectors: pCMV-TO/FRT-Emerald (GFP); pcDNA5-TO/FRT-mCherry (only LUZP1); pcDNA5-TO/FRT-FLAG; pPUR 5’/FRT/TO-FLAG-BirA* (only LUZP1). The GFP-LUZP1, GFP-EPLIN and mCherry-LUZP1 fusions’ coding sequences were amplified by PCR and sub-cloned into the lentiviral vector pHR-SIN-SFFV. The LUZP1 truncation mutants were generated by amplifying the respective coding sequences by PCR, followed by ligation into the pCMV-TO/FRT-Emerald (GFP) or pcDNA5-TO/FRT-FLAG vectors.

### Lentiviral production and generation of the hTERT RPE-1 stable cell lines

For the production of lentiviral particles, HEK293T cells were co-transfected with pHR-SIN-SFFV-GFP-LUZP1, pHR-SIN-SFFV-mCherry-LUZP1, pHR-SIN-SFFV-GFP-EPLINα, or pHR-SIN-SFFV-GFP-EPLINβ, and the second generation packaging (pCMV-dR8.74psPAX2) and envelope (pMD2.G) plasmids, using the Lipofectamine® 3000 transfection reagent (Invitrogen) according to the manufacturer’s instructions. Lentiviral particles in conditioned media from HEK 293T cells, collected at 48h post transfection, were used to transduce hTERT RPE-1 cells. GFP or GFP + mCherry positive cells were cell sorted to establish the final fluorescent cell lines.

### Treatment with cytoskeleton drugs

To depolymerize microtubules, hTERT RPE-1 GFP-LUZP1 cells were incubated with 30µM nococazole for 1h at 37°C before they were fixed and processed for immunofluorescence. To perturb the actin cytoskeleton, cells were treated with 50nM cytochalasin D for 16h. In the case of RNAi experiments, the cytochalasin treatment happened during the last 16h of the 72h of gene silencing. For all experiments, the cells were treated with the same volume of DMSO as a control.

### Generation and characterization of stable and inducible HEK293 pools for BioID

Flp-In™ T-REx™ HEK293 cells were co-transfected with pOG44 (Flp-recombinase expression vector) and the pPUR 5’/FRT/TO-FLAG-BirA*-LUZP1. Transfections were performed with Lipofectamine® 2000 according to manufacturer’s instructions. After transfection, cells were selected with 1μg/ml puromycin. Cells were incubated in one of three conditions for 24h: 1) 1μg/mL tetracycline (Bioshop) and 50 μM biotin (Sigma); (2) 1 μg/ml tetracycline only; (3) no tetracycline or biotin added. The cells were fixed with ice-cold methanol and processed for immunofluorescence. Biotinylated proteins were detected using fluorophore-conjugated streptavidin (Invitrogen).

### BioID sample preparation

BioID (Roux et al., 2012) was carried out essentially as described previously (Coyaud et al., 2015). Independent replicates of five 15 cm plates of sub-confluent (60%) stably expressing FLAG-BirA* alone (eight replicates) or FLAGBirA*-LUZP1 (2 replicates) were incubated for 24h in complete media (and in serum free media for 48h prior to biotin addition for the ciliated condition) supplemented with 1 µg/ml tetracycline (Sigma) and 50 µM biotin (BioShop). Cells were collected and pelleted (2,000 rpm, 3 min), the pellet was washed twice with PBS and dried pellets were snap frozen. The cell pellet was resuspended in 10 ml of lysis buffer (50 mM Tris-HCl pH 7.5, 150 mM NaCl, 1 mM EDTA, 1 mM EGTA, 1% Triton X-100, 0.1% SDS, 1:500 protease inhibitor cocktail (Sigma-Aldrich), 1:1,000 benzonase nuclease (Novagen) and incubated on an end-over-end rotator at 4°C for 1 h, briefly sonicated to disrupt any visible aggregates, then centrifuged at 45,000 x g for 30 min at 4°C. Supernatant was transferred to a fresh 15 mL conical tube. 30 μl of packed, pre-equilibrated Streptavidin sepharose beads (GE) were added and the mixture incubated for 3 hours at 4°C with end-over-end rotation. Beads were pelleted by centrifugation at 2,000 rpm for 2 min and transferred with 1 mL of lysis buffer to a fresh Eppendorf tube. Beads were washed once with 1 mL lysis buffer and twice with 1 mL of 50 mM ammonium bicarbonate (pH=8.3). Beads were transferred in ammonium bicarbonate to a fresh centrifuge tube and washed two more times with 1 ml ammonium bicarbonate buffer. Tryptic digestion was performed by incubating the beads with 1 µg MS-grade TPCK trypsin (Promega, Madison, WI) dissolved in 200 μl of 50 mM ammonium bicarbonate (pH 8.3) overnight at 37°C. The following morning, 0.5 μg MS-grade TPCK trypsin was added, and beads were incubated 2 additional hours at 37°C. Beads were pelleted by centrifugation at 2,000 x g for 2 min, and the supernatant was transferred to a fresh Eppendorf tube. Beads were washed twice with 150 µL of 50 mM ammonium bicarbonate, and these washes were pooled with the first eluate. The sample was lyophilized and resuspended in buffer A (0.1% formic acid). 1/5th of the sample was analyzed per MS run.

### Mass spectrometry

Analytical columns (75-um inner diameter) and pre-columns (150-um inner diameter) were made in-house from fused silica capillary tubing from InnovaQuartz (Phoenix, AZ) and packed with 100 Å C18–coated silica particles (Magic, Michrom Bioresources, Auburn, CA). Peptides were subjected to liquid chromatography (LC)-electrospray ionization-tandem mass spectrometry, using a 120 min reversed-phase (100% water–100% acetonitrile, 0.1% formic acid) buffer gradient running at 250 nl/min on a Proxeon EASY-nLC pump in-line with a hybrid LTQ-Orbitrap Velos mass spectrometer (Thermo Fisher Scientific, Waltham, MA). A parent ion scan was performed in the Orbitrap using a resolving power of 60,000, then up to the twenty most intense peaks were selected for MS/MS (minimum ion count of 1000 for activation), using standard collision induced dissociation fragmentation. Fragment ions were detected in the LTQ. Dynamic exclusion was activated such that MS/MS of the same m/z (within a range of 15 ppm; exclusion list size = 500) detected twice within 15 s were excluded from analysis for 30 s. For protein identification, Thermo .RAW files were converted to the .mzXML format using Proteowizard (Kessner et al., 2008), then searched using X!Tandem (Craig and Beavis, 2004) and COMET (Eng et al., 2013) against the human Human RefSeq Version 45 database (containing 36,113 entries). Data were analyzed using the trans-proteomic pipeline (TPP) (Deutsch et al., 2010; Pedrioli, 2010) via the ProHits software suite (v3.3) (Liu et al., 2010). Search parameters specified a parent ion mass tolerance of 15 ppm, and an MS/MS fragment ion tolerance of 0.4 Da, with up to 2 missed cleavages allowed for trypsin. Variable modifications of +16@M and W, +32@M and W, +42@N-terminus, and +1@N and Q were allowed. Proteins identified with an iProphet cut-off of 0.9 (corresponding to ≤1% FDR) and at least two unique peptides were analyzed with SAINT Express v.3.3.1. Eight control runs (from cells expressing the Flag-BirA* epitope tag) were collapsed to the two highest spectral counts for each prey and compared to the two biological replicates of *LUZP1* BioID (two technical replicates of each; four runs collapsed to three highest spectral counts for each prey) in normal and serum starved conditions. High confidence interactors were defined as those with Bayesian false discovery rate (BFDR) ≤0.01.

### RNA interference

To silence LUZP1 and EPLIN, RPE□1 cells (1 × 10^5^ cells seeded in 6□well plates) were transfected with 40nM (final concentration) of a pool of 2 siRNAs targeting either gene obtained from Dharmacon (ON□TARGET plus) using the Lipofectamine® RNAiMAX transfection reagent (Invitrogen) according to the manufacturer’s instructions. The Luciferase GL2 Duplex non-targeting siRNA from Dharmacon was used as a negative control. Gene silencing was carried out for 72h. The siRNA sequences are listed in supplementary table 2. When indicated, the cells were serum-starved (DMEM/F12 supplemented with GlutaMAX™ and sodium bicarbonate (1.2 g/l) but without serum) for 72h to induce the formation of primary cilia.

### RNAi rescue experiments

hTERT RPE-1 GFP-only and hTERT RPE-1 GFP-LUZP1 stable lines expressing the siRNA resistant or sensitive LUZP1 transgenes were transfected with 40 nM (final concentration) of a pool of 2 siRNAs (Dharmacon ON□TARGET plus) directed against LUZP1 using the Lipofectamine® RNAiMAX transfection reagent (Invitrogen) according to the manufacturer’s instructions. The Luciferase GL2 Duplex non-targeting siRNA from Dharmacon was used as a negative control. Gene silencing was carried out for 72h. hTERT RPE-1 GFP-only and hTERT RPE-1 GFP-EPLINα stable lines were transfected with 40 nM (final concentration) of a pool of 2 siRNAs (Dharmacon ON□TARGET plus) directed against the 3’UTR of EPLIN transcripts. Gene silencing was carried out for 72h.

### Fluorescence microscopy

For immunofluorescence, the cells were fixed with cold methanol (10 min at −20°C), blocked with 0.2% Fish Skin Gelatin (Sigma-Aldrich) in 1x PBS (20 min), incubated with the primary antibodies in blocking solution (1h), washed with blocking solution and incubated with fluorophore-conjugated secondary antibodies (Molecular Probes) and DAPI in blocking solution (1h). After a final wash in blocking solution the coverslips were mounted on glass slides by inverting them onto mounting solution (ProLong Gold antifade, Molecular Probes). For the characterization of the HEK293 Flag-BirA*-LUZP1 line, the cells were also incubated with fluorophore-conjugated streptavidin (Molecular Probes). The cells were imaged on a DeltaVision (Applied Precision) imaging system equipped with an IX71 microscope (Olympus), CCD camera (CoolSNAP HQ2 1024×1024, Roper Scientific) and a ×60/1.42 NA planApochromat oil-immersion objective (Olympus). Z stacks (0.2□μm apart) were collected, deconvolved using softWoRx (v5.0, Applied Precision) and are shown as maximum intensity projections (pixel size 0.1064□μm). Primary and secondary antibodies used are listed in supplementary table 2.

For live-imaging, the cells were seeded in Nunc™ Lab-Tek™ Chamber Slides and imaged on the DeltaVision system with temperature and CO_2_ control, using a x40/1.35 NA oil-immersion objective (Olympus). Time lapse was 5 min. Z stacks (1□μm apart) were collected, deconvolved using softWoRx (v5.0, Applied Precision) and are shown as maximum intensity projections (pixel 0.33463□μm). Images were analysed, including cilia measuring, with the FIJI (ImageJ; National Institutes of Health) software.

### Cell proliferation assay

The time-lapse imaging of siRNA-transfected cells was done using an IncuCyte® S3 Live Cell Analysis System (Essen BioScience) using a 10x objective. Phase images were acquired every 3h for 72h in 5% CO_2_ humidified atmosphere at 37ºC. Cell proliferation was quantified using the IncuCyte® S3 Cell by Cell analysis module.

### Western Blotting

For Western blots, the cells were collected, lysed in Laemmli buffer and treated with benzonase nuclease (Sigma). Proteins were separated by loading whole cell lysates onto an 8%-10% SDS-PAGE gel for electrophoresis and then transferred to a PVDF membrane (Immobilon-P, Millipore). Membranes were incubated with primary antibodies in TBST (TBS, 0.1% Tween-20) in 5% skim milk powder (Bioshop), supplemented with 2.5% BSA Fraction V (OmniPur) in the case of FLAG western blots. Blots were washed 3x 10 min in TBST, then incubated with secondary HRP-conjugated antibodies. Western blots were developed using SuperSignal reagents from Thermo Scientific.

### Co-Immunoprecipitation followed by western blot

For co-immunoprecipitation of Flag fusions, the respective HEK293 stable lines were incubated with tetracycline (1µg/ml) for 24h after which they were washed with 1x PBS, harvested and frozen at −80ºC or lysed immediately (50mM HEPES pH8; 100mM KCl; 2mM EDTA; 10% Glycerol; 0.1% NP-40; 1mM DTT; protease inhibitors) for 30 min on ice. The lysates were then frozen in dry ice for 5 min and then thawed and centrifuged for 20 min at 16,000xg at 4ºC. The cleared lysates were then incubated with ANTI-FLAG^®^ M2 Affinity Gel (Sigma) over-night at 4ºC. A fraction of the protein extracts (Inputs) were saved before the incubation with the beads. After the incubation, the beads were pelleted and washed with lysis buffer. The samples (Inputs and IPs) were prepared for SDS-PAGE by addition of Laemmli buffer and boiling. The proteins were transferred to PVDF membranes (Immobilon-P, Millipore) and probed with antibodies to detect the FLAG fusions and endogenous proteins. The GFP were performed similarly using extracts from HEK293 or RPE-1 cells expression GFP-fusions. Protein G Sepharose 4 Fast Flow (P3296 Sigma-Aldrich) were incubated with 2 µg of GFP antibody raised in Goat for 2h at 4°C and were then washed with lysis buffer. GFP antibody-conjugated beads were then used to pull-down the GFP-fusions. Primary and secondary antibodies used are listed in supplementary table 2.

### Statistical methods

All p-values are from two-tailed unpaired Student t-tests. All error bars are S.D. Individual p-values, experiment sample numbers and the number of replicates used for statistical testing are reported in corresponding figure legends. ***p<0.001, **p<0.01, *p<0.05. NS – non significant.

## AUTHOR CONTRIBUTIONS

J.G. conceived the project, designed the research plan, and performed the experiments and data analyses. Sample prep for MS was carried out by E.M.N.L., E.C. and B.R. performed the MS and the related analyses. J.G. wrote the manuscript with contributions from the other authors. L.P. supervised and funded the project.

## AKNOWLEDGEMENTS

We are grateful to Dr. Johnny Tkach, Dr. Ladan Gheiratmand, and Dr. Mikhail Bashkurov for very fruitful discussions during the development of this work. J.G. was partially funded by a post-doctoral fellowship from Fundação para a Ciência e a Tecnologia (SFRH/BPD/75847/2011). This work was funded from operating grants from the CIHR (MOP 142492) and the Krembil Foundation to L.P

